# Rethinking GPS Navigation: Creating Cognitive Maps Through Auditory Clues

**DOI:** 10.1101/2020.05.13.094219

**Authors:** Gregory D. Clemenson, Antonella Maselli, Alex Fiannaca, Amos Miller, Mar Gonzalez-Franco

**Affiliations:** Microsoft Research, Redmond, Washington 98052, USA; Department of Neurobiology and Behavior, University of California Irvine, Irvine, California 92697, USA

**Keywords:** GPS, mental map, navigation, hippocampus

## Abstract

GPS navigation is commonplace in everyday life. While it has the capacity to make our lives easier, it is often used to automate functions that were once exclusively performed by our brain. Staying mentally active is key to healthy brain aging. Therefore, is GPS navigation causing more harm than good? Here we demonstrate that traditional turn-by-turn navigation promotes passive spatial navigation and ultimately, poor spatial learning of the surrounding environment. We propose an alternative form of GPS navigation based on sensory augmentation, that has the potential to fundamentally alter the way we navigate with GPS. By implementing a 3D spatial audio system similar to an auditory compass, users are directed towards their destination without explicit directions. Rather than being led passively through verbal directions, users are encouraged to take an active role in their own spatial navigation, leading to more accurate cognitive maps of space. Technology will always play a significant role in everyday life; however, it is important that we actively engage with the world around us. By simply rethinking the way we interact with GPS navigation, we can engage users in their own spatial navigation, leading to a better spatial understanding of the explored environment.

## Introduction

Navigation is a fundamental human behavior that is critical for everyday life. Exploring new environments, understanding the current location, and remembering how to get back are essential for successful navigation (1). In fact, knowing where to go and where we came from are such fundamental processes that they were once critical to our own ancestral survival (2). But with the arrival of GPS and navigation apps, is this a thing of the past? And more importantly, if these apps are now responsible for a function that is performed by our brain and was once critical to our survival, is it doing more harm than good?

Navigation requires the coordination of numerous perceptual and sensory processes that are further supported by several brain regions, including the hippocampus, retrosplenial cortex, striatum and entorhinal cortex, to create successful representations of space (3–8). While spatial cognition is not solely attributed to one brain region, the hippocampus is well known to play a critical role in spatial learning, memory and navigation, and has long been thought to contain the “cognitive map”(8, 9). In fact, the creation of cognitive maps, development of spatial expertise, and exploration of novel environments can have a positive impact on hippocampal structure and function (10–15). As we age, however, our hippocampus and spatial abilities decline (6, 16–21), even predicting the conversion from mild cognitive impairment to Alzheimer’s disease (22–24). Therefore, creating mental maps is not only important for the navigation of daily life, but is critically important as we age.

At the most fundamental level, cognitive maps are formed through exploration (8, 9, 25–28). While the animal work has described the formation of these spatial neural networks in detail, studies have demonstrated that humans are quite capable of learning about space in other ways such as with the use of (paper) maps (29, 30). Most significantly, the literature describes two main types of navigation strategies: egocentric and allocentric. Egocentric navigation is considered striatal and describes how cues within the environment relate to the individual (a set of directions) whereas allocentric navigation is hippocampal and describes how cues within the environment relate to one another (a map). While both navigation strategies have their advantages, they are thought to operate both independently and in parallel (31). A true cognitive map requires an allocentric perspective of space that the hippocampus provides, so that the spatial information is flexible and can be used from any location (9). Furthermore, a successful creation of cognitive maps requires active engagement in the navigation process as spatial decision making is the primary component of active learning for the acquisition of map based knowledge (3, 32).

Is it possible to find a balance between our internal navigation system and modern technology? Certainly, new technology has allowed us to go further and reach unexplored places in ways we would have only imagined prior to GPS access on our mobile phones. One can get lost in the streets of Tokyo and still find their way back without fear. Hence, we have reached a modern paradox for navigation: navigation apps allow us to explore more places, while at the same time making us worse explorers. Here, we argue that current GPS apps (based on turn-by-turn navigation) promote a passive form of navigation that does not support learning or the formation of cognitive maps (33). Turn-by-turn navigation, is a passive navigation system based on an egocentric perspective. The user does not make any decisions about the navigation process, they simply input a desired location and follow the directions on the app.

While traditional turn-by-turn navigation is effective in its ability to lead us to a desired location, this passive form of navigation does not support spatial learning, thus having a detrimental impact on humans navigation skills and spatial cognition. Several studies have observed a link between GPS navigation and poor spatial awareness. Anthropologists studying Inuit hunters in northern Canada have noted their extraordinary ability to navigate the arctic tundra, despite the harsh, desolate terrain and the lack of reliable landmarks across large spaces. While the use of traditional navigation tools, like maps and compasses, has empowered their extraordinary perceptual skills, that were developed and passed down from one generation to the next; GPS technologies replaced the need for such skills altogether. Younger generations of Inuit hunters have traded in these deep-rooted wayfinding skills for modern GPS technology. While easy to use, GPS technology could not help the young Inuit hunters adapt to the extreme weather or hazards of the arctic tundra. An over dependence on GPS technology led to an increase in serious accidents and death, as these young Inuit hunters were essentially navigating “blindfolded” (2). Even in modern cities, in-car GPS navigation “inhibits the process of experiencing the physical world” (33). Rather than aiding our ability to navigate, GPS navigation is designed to relieve us of our involvement and disengage us from our surrounding environment. After all, GPS navigation is a form of destination-oriented transport in which humans become a “passenger of their own body”, as opposed to wayfaring, which embeds us into the environment as the driver, in order to find the destination (34).

Here we suggest an alternative form of GPS navigation, that is possible with the use of 3D spatialized audio. We position an auditory beacon, which is essentially a continuous sound virtually positioned in 3D, to always be emanating from the direction straight from the destination it is associated with. By the means of this auditory beacon, users of GPS apps can regain their active role, and navigate in the space like they used to: without delegating decisions (see materials section for more details about the beacon implementation). Users navigate in the direction from which the beacon sounds, constrained only by the available paths traveling in that direction. In this scenario, the mobile app acts as a compass that directs the user towards the destination in an allocentric manner. The goal of our research is to demonstrate that by simply rethinking the way we interact with technology and introducing sensory augmentation into the equation (in the form of 3D audio) we can have a real impact on modern navigation without having to compromise our own internal navigation system or our ability to create mental maps.

## Materials and Methods

### Participants

In total, we recruited 53 (19 female, 34 male; mean age = 33.8, SD = 8.21) individuals to participate in the study. Using a Microsoft email blast, we recruited both Microsoft Employees (experts) and Interns (naïve). All participants gave informed consent and were compensated with a $50 Amazon gift card upon completion of the study. The study was conducted in accordance with Microsoft IRB guidelines.

### Experimental Groups

Participants were classified as either Experts or Naïve based on their experience with the Microsoft campus. The Naïve group (n=24, 9 female, 15 male; mean age = 29.9, SD = 7)consisted on Summer Interns at Microsoft who had just begun their internship. All participants in the Naïve group had been on the Microsoft campus for less than three weeks at the time of the study. The Expert group (n=29, 10 female, 19 male; mean age = 37.79, SD = 8.2) consisted of Microsoft employees who had been on the Microsoft campus for at least six months at the time of the study (up to +20 years). Within each group, individuals were randomly divided into two conditions: Turn-by-turn and Beacon. The Turn-by-turn group performed the task using turn-by-turn directions and the Beacon group performed the task using the Microsoft Soundscape app^1^.

Prior to the study, participants were asked two questions about their familiarity with the Microsoft campus and their navigation abilities (Seven-point Likert scale for each question). These self-reports of familiarity with the Microsoft campus and confidence in their own navigation abilities confirmed that the Naïve group was less familiar with the campus (Naïve: mean = 2.1, SD = 0.92; Experts: mean = 4.7, SD = 1.44, p=0.04) but not significantly less confident in their own navigation (Naïve: mean = 3.9, SD = 1.45; Experts: mean = 4.3, SD = 1.78) compared to the Expert group (p=0.4).

### Beacon Implementation

The audio beacon consists of continuously playing audio that is localized around the user such that the source always appears to be coming from the direction towards the desired destination. The rendering of the audio stream is performed using a head-related transfer function (43), and the audio stream is localized one meter away from the user in the direction towards the desired location (Figure 4).

**Figure 4.**
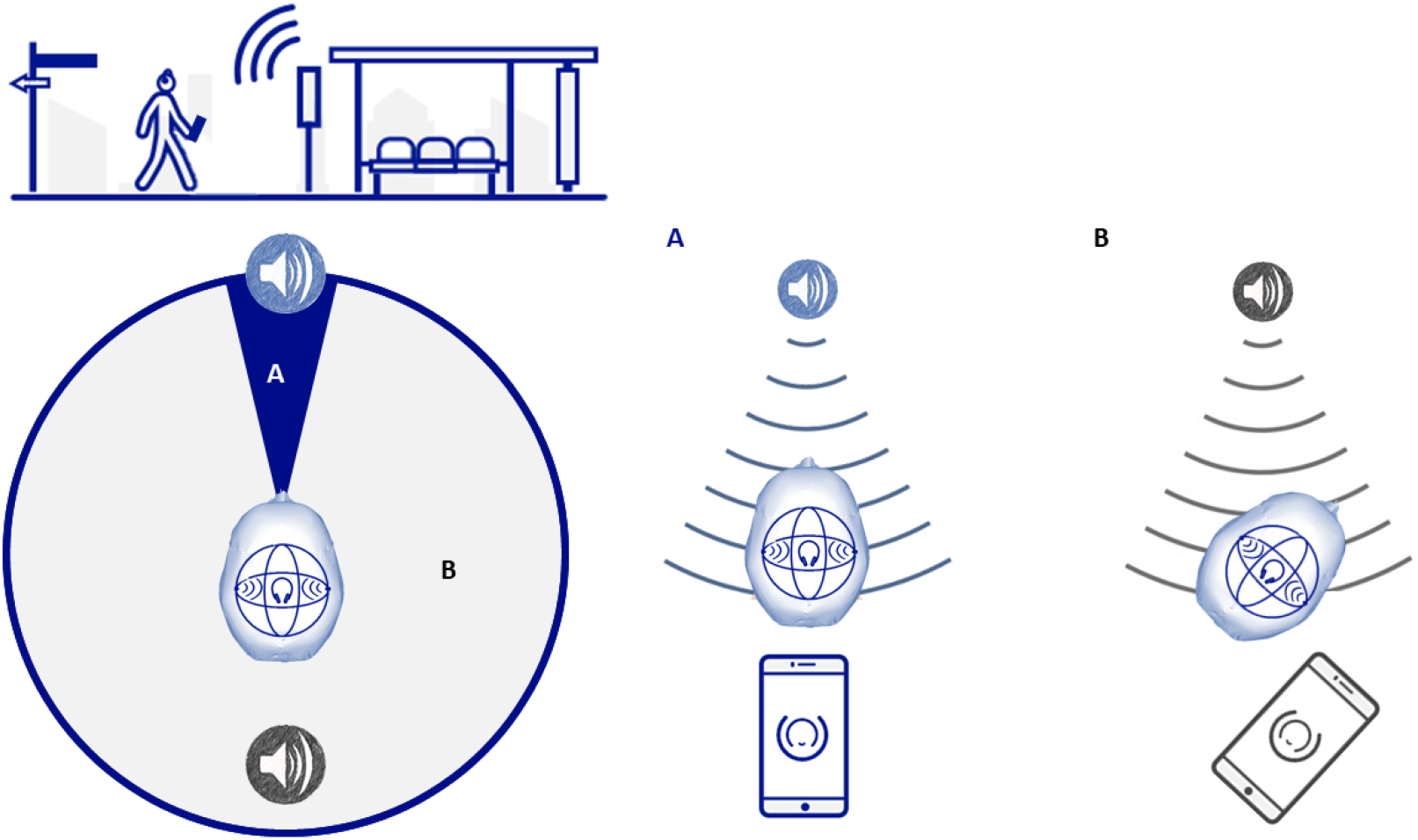
Implementation of the Beacon. The participant receives spatial audio information as coming from the beacon source. To amplify the information of directionality, we further implement two types of sounds depending on whether the participant is pointing directly at the beacon (A) or it is walking in another direction (B). This small design decision guides participants into finding the beacon faster.

The localization of the audio is updated continuously based on the user’s orientation (as determined by the gyroscope and magnetometer in the user’s phone running Soundscape, Figure 4). The beacon’s audio varies based on the angle between the user’s orientation and the bearing from the user’s location to the location of the beacon’s destination. If the angle is greater than 22.5°, a base audio track consisting of a clip-clop sound is played. If the angle is less than 22.5°, the same base audio Is played, but an additional audio track consisting of a high pitch ting sound synchronized with the clip-clop sound is added in. This additional layer of audio provides a secondary encoding of the directionality of the beacon. It is important to note that the audio was designed to have a minimal footprint into the auditory space of the participant so it would not occlude the understanding of the real environment, without affecting sights and sounds of the world around you (44).

We believe that auditory beacons will fall on the category of navigation aids, and are not designed to directly substitute human wayfinding abilities, but to augment the perceptual navigation (1,44). Different beacon configurations have been explored for that purpose (46–48). One of the earliest implementations from 2002 AudioGPS (46) showed positive informal comments towards audio guided vector direction navigation. Later gpsTunes and Ontrack worked on a similar way but based on the music that the user was listening (47, 48). Audio Bubbles tested whether this type of tools could be used for tourist wayfinding (49). However, none of the previous efforts carried a formal evaluation of the mental map creation with these tools.

A clip of the audio is provided in the supplementary material, but we encourage to download the Microsoft Soundscape app for a full experience and demo of the beacon implementation.

### Soundscape Scavenger Hunt

Soundscape is a spatial awareness app, original designed to assist seeing-impaired individuals in learning about the environment around them. Soundscape uses a combination of features to both guide users to a desired location (audio beacon) and to inform users about nearby places of interest or intersections (callouts). For the purposes of this experiment, we disabled the callouts feature of the app to ensure that participants were navigating using only the audio beacons (continuous 3D spatialized audio that always emanates from the direction towards the desired location).

All participants participated in a scavenger hunt activity in which they had to locate five different POIs, of varying degrees of difficulty to find, on the Microsoft campus (Figure 1). There were some POIs we expected some users to have seen before (volleyball court), but others that were not immediately identifiable (two benches and hidden door or a particular building).

**Figure 1.**
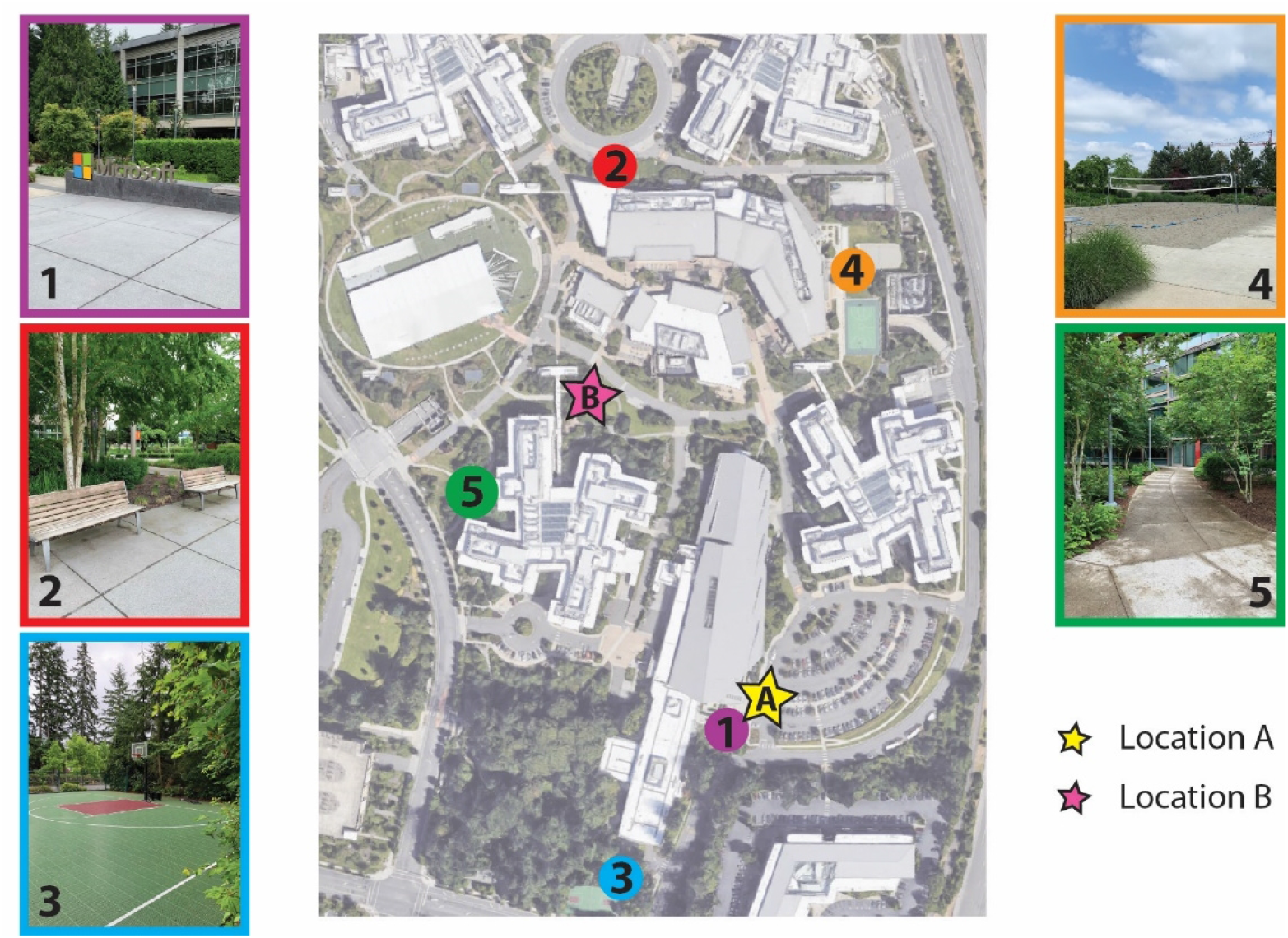
Overhead view of the scavenger hunt on the Microsoft campus. POI locations are denoted by the numbers and colored circles: Microsoft Sign (1; purple), Two Benches (2; red), Basketball Court (3; blue), Volleyball Court (4; orange), and Hidden Door (5; green). The locations of the pointing tasks are denoted with stars: Location A (A; yellow) and Location B (B; pink).

The Microsoft campus is very large, spanning over 2 square kilometers, however the selected area was relatively central, and we reduced the scavenger hunt to a 0.12 square kilometers region.

The order of the POIs was picked to purposely avoid a direct path from one to the next, this way participants had to make active decisions on where to go. Pictures of each POI were displayed in the Soundscape app, one at a time, and participants were instructed to find the pictured location in the real world. See supplementary material (Supplementary Figure 3A and 3B), for further details on the scavenger hunt visualization in the app. While both the Turn-by-turn and Beacon groups used Soundscape to view the POIs, the Turn-by-turn group navigated to the POIs through verbal turn-by-turn directions (by the experimenter; Supplementary Figure 4) and the Beacon group navigated to the POIs through the use of audio beacons in Soundscape. The Turn-by-turn received verbal directions (directly taken from a navigation app) from the experimenter because turn-by-turn directions did not have the spatial resolution to determine the precise location of each turn juncture on the Microsoft campus. Neither the Turn-by-turn group or the Beacon group were allowed to view their location on a visual map. Therefore, this feature was removed from the current study as the goal was to test the effects of auditory stimulation. In addition, a visual map could be more time consuming that with the use of an auditory navigation system (49). The Beacon group was not allowed to make shortcuts through buildings. Once the POI was found (determined by Soundscape based on GPS location), a picture of the next POI was presented, the audio beacon was updated to point towards the new POI’s location, and participants were instructed to continue to the next POI. The presentation order of the POIs was the same for all participants and was specifically arranged so that every POI had multiple access routes.

Soundscape recorded timestamps, GPS locations and heading directions at a sampling rate of approximately 1Hz throughout the entire duration of the study.

### Experimental Design

Two tasks were used to assess spatial knowledge. The first was a pointing task, which is commonly used to assess spatial abilities in real-world studies (50–52). In the pointing task (Supplementary Figure 3), a picture of a POI was presented on the phone and participants had to point, using the phone, from their current location to the POI. To confirm the direction, participants pressed a button on the screen and the Soundscape app recorded the direction. As there is inherent error in the mobile phone compass, we used relative differences between heading directions when computing the amount of pointing error as opposed to comparing the angle differences between true direction and the heading direction provided by the phone. For example, the first POI (Microsoft Sign) is visible from Location A (Figure 1) and was therefore used as the reference point for all other POIs. In other words, the amount of error for each POI is the angle difference between the POI and the reference POI (Microsoft Sign), as provided by the phone, versus the angle difference between the true direction of the POI and the true direction of the reference POI (Microsoft Sign). In Location B, however, we did not have a reference POI visible. To determine the pointing error for each POI, we calculated the average error between the current POI and all other POIs, used as a reference point. For example, pointing error for the Microsoft Sign is the average of the angle difference between the Microsoft sign and Two Benches, Microsoft Sign and Basketball Court, Microsoft Sign and Volleyball Court, and Microsoft Sign and Hidden Door. True direction was determined using GPS coordinates of the current location and the POI.

In the second task, participants were given a simple map of the Microsoft campus (with no distinguishing features or labels) and asked to mark the locations of all five POIs (Supplementary Figure 5). Participant maps were scanned and overlaid with a schematic containing the actual POI locations, as determined by GPS coordinates. ImageJ (imagej.nih.gov) was used to analyze the distance (in pixels and later converted into meters) between the estimated location and the real location of all POIs, to come up with an error metric.

Prior to the scavenger hunt, participants performed a pre-test pointing task at the start location (Location A) to determine familiarity with each of the POIs. After completing the scavenger hunt, participants performed two post-test pointing tasks, one from the same location as the pre-test (location A in Figure 1) and one from an entirely novel location (location B in Figure 1). Lastly, all participants were asked to draw the POIs on the map given.

All participants were given the same instructions and were naïve to the details of the study. They were told that they are participating in a scavenger hunt in which they must find five different locations spread throughout the campus.

The Beacon group was taught how to use/interpret the 3D sound with verbal directions and the very first landmark (Figure 1: Landmark 1). Standing at Location A, participants were taught what a beacon sounded like and how to interpret the direction. The Turn-by-turn groups was also given instructions for how to follow turn-by-turn directions based on the very first landmark (Figure 1: Landmark 1). At every landmark they were told that this was their N location and then started going for the next landmark immediately.

### Statistical Analyses

All statistics were performed using Prism 8 (www.graphpad.com) and specific statistical tests used are displayed next to each result.

## Results

We prepared a scavenger hunt with five different Points of Interest (POIs) that participants (n=53) had to visit using either auditory beacons or turn-by-turn directions (Figure 1). Participants were either experts in the terrain (with over 6 months of exposure) or completely naïve (who did not know the area, had been there for a maximum of 3 weeks).

### There were no differences between groups or conditions in familiarity with the POIs prior to the scavenger hunt

Prior to starting the scavenger hunt, we determined a baseline familiarity with the POIs. All participants stood in the same spot and performed a pointing task in which they pointed to all five POI locations, even though they were not visible. Using the amount of error as a measure of accuracy, we performed a 2×2 ANOVA across group (Expert and Naïve) and condition (Turn-by-turn and Beacon). We did not find a significant main effect of group (Figure 2A; F(1,49) = 0.55, p = 0.46), condition (F(1,49) = 0.003, p = 0.96), or interaction (F(1,49) = 0.09, p = 0.76), showing that at pre-test, there were no differences between groups in their ability to point to each of the POI locations (All groups: Mean = 48.55 degrees, SD = 19.53). See supplementary material for individual POI performance (Supplementary Figure 1A and 1B).

**Figure 2.**
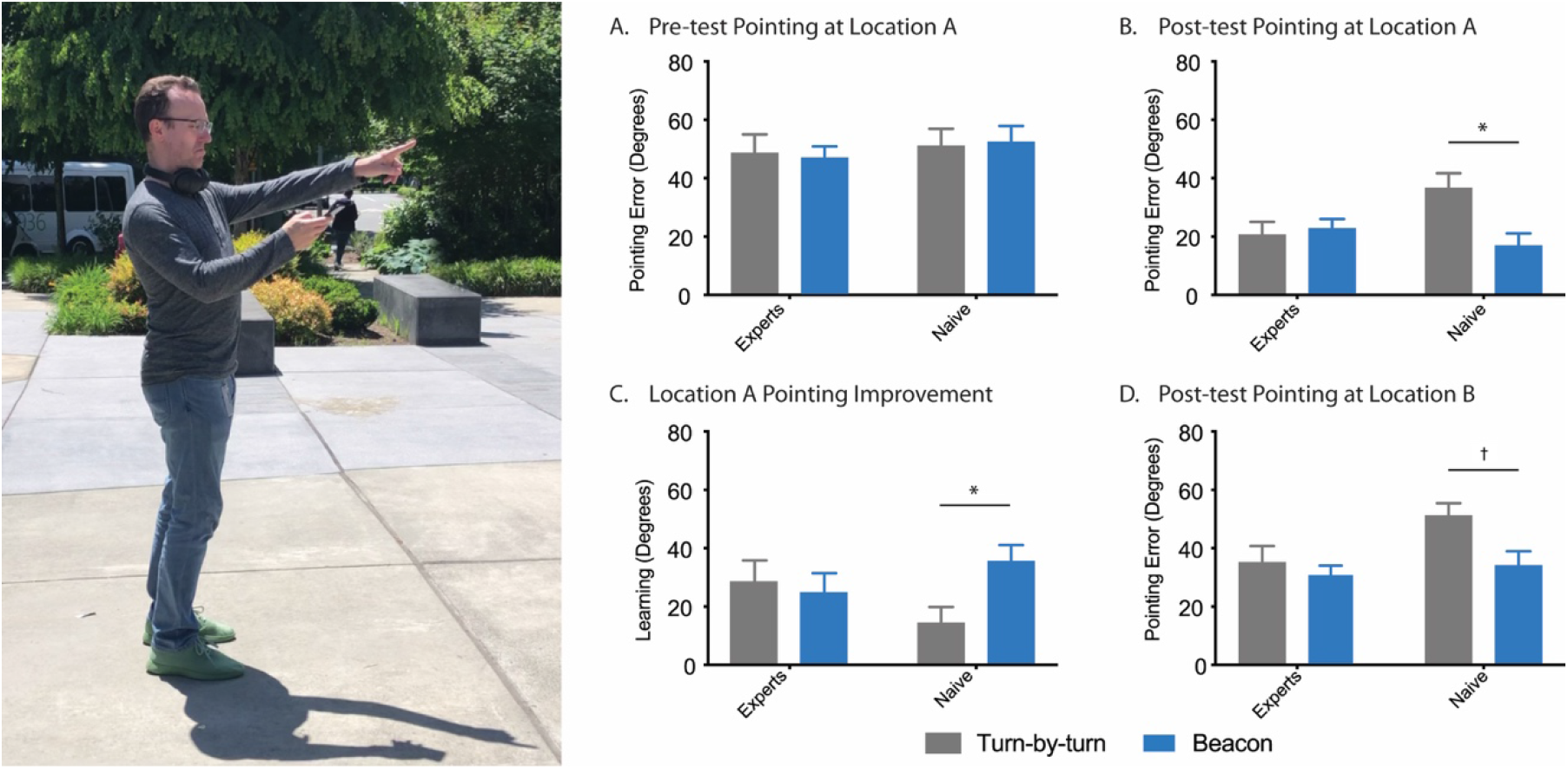
Performance of all groups and conditions on the pre-test and post-test pointing tasks. (A) Pointing error of all groups and conditions from Location A at pre-test. (B) Pointing error of all groups and conditions from Location A at post-test. (C) Improvement of all groups and conditions when pointing from Location A (pre-test minus post-test pointing error). (D) Pointing error of all groups and conditions from novel Location B at post-test. All data are presented as ± SEM, * p < 0.05, † p < 0.1

### Upon completion of the scavenger hunt, participants in the Beacon condition were more accurate at pointing from the initial start location (A)

To determine how well participants learned the locations of the visited POIs, we performed a pointing task, after the scavenger hunt, from the same starting position as the initial pre-test pointing task (Location A). A 2×2 ANOVA across group (Expert and Naïve) and condition (Turn-by-turn and Beacon), demonstrated a significant interaction (Figure 2B; F(1,49) = 6.2, p < 0.05) and main effect of condition (F(1,49) = 4.41, p < 0.05), but no main effect of group (F(1,49) = 1.53, p = 0.22). A post-hoc analysis (Tukey’s correction for multiple comparisons) revealed that the Naïve Beacon group (Mean = 16.9 degrees, SD = 13.98) was significantly better than the Naïve Turn-by-turn group (Mean = 36.64 degrees, SD = 18.1). These data indicate that the Beacon group, especially Naïve Beacon users, displayed a better knowledge of the POI locations. See supplementary material for individual POI performance (Supplementary Figure 1C and 1D).

### The Naïve Beacon group demonstrated a significant improvement in learning, from pre-test to post-test

To assess the amount of learning that occurred from pre-test to post-test, we compared performance on the pointing task from location A at post-test and pre-test, using the difference in accuracy (post-test pointing error minus pre-test pointing error) as a metric of learning. A 2×2 ANOVA across group (Expert and Naïve) and condition (Turn-by-turn and Beacon) revealed a significant interaction (Figure 2C; F(1,49) = 3.96, p = 0.05) but no main effect of group (F(1,49) = 0.03, p = 0.85) or condition (F(1,49) = 1.96, p = 0.16). While a post-hoc analysis (Tukey’s correction for multiple comparisons) did not reveal any significant differences in learning between groups, it is worth noting that a simple t-test demonstrated a difference between the Naïve Beacon (Mean = 35.55 degrees, SD = 18.36) and Naïve Turn-by-turn groups (Mean = 14.42 degrees, SD = 19.83). See supplementary material for individual POI performance (Supplementary Figure 1E and 1F).

### Auditory navigation improved pointing accuracy from a novel location (location B)

In addition to pointing from the pre-test location, participants also pointed to all POIs from an entirely novel location (Figure 1, Location B) to assess how well they acquired an allocentric representation of space. A 2×2 ANOVA across group (Expert and Naïve) and condition (Turn-by-turn and Beacon) revealed a significant main effect of both group (F(1,49) = 4.05, p < 0.05) and condition (Figure 2D; F(1,49) = 5.01, p < 0.05) but not interaction (F(1,49) = 1.96, p = 0.17). After correcting for multiple comparisons (Tukey) there was a trend towards a difference between the Naïve Turn-by-turn (Mean = 51.18 degrees, SD = 15.37) and Naïve Beacon group (Mean = 34.12 degrees, SD = 16.05), however this was not significant (p = 0.07). It is worth noting that, a simple t-test demonstrated a difference between the Naïve Turn-by-turn and Naïve Beacon groups (t-test, p = 0.01). See supplementary material for individual POI performance (Supplementary Figure 2A and 2B).

### The Beacon groups were better at identifying the POI locations on a drawn map

After the scavenger hunt, participants were asked to identify the POI locations on a simple map outline of the area with no labels (Figure 3A). A 2×2 ANOVA across group (Expert and Naïve) and condition (Turn-by-turn and Beacon) revealed a significant main effect of condition (Figure 3B; F(1,49) = 7.3, p < 0.01) but not group (F(1,49) = 2.7, p < 0.1) or interaction (F(1,49) = 3.0, p = 0.08). A post-hoc analysis (Tukey’s correction for multiple comparisons) revealed a significant difference between the Naïve Turn-by-turn (Mean = 177.55 meters, SD = 97.0) and Naïve Beacon (Mean = 86.1 meters, SD = 64.51) groups (p < 0.05). See supplementary material for individual POI performance (Supplementary Figure 2C and 2D).

**Figure 3.**
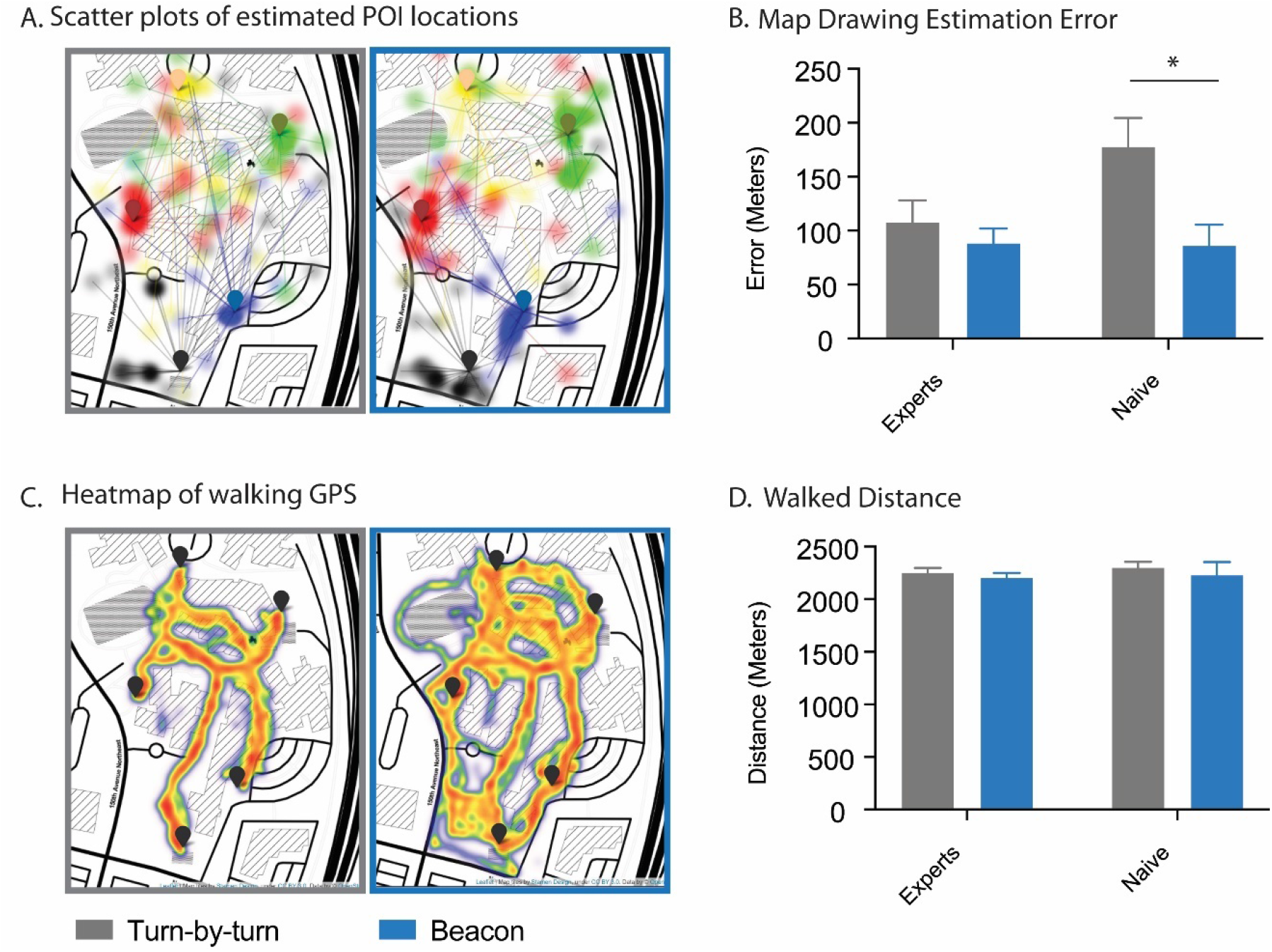
POI accuracy, path distance and routes taken during the scavenger hunt. (A) Estimated locations of all POIs of the Turn-by-turn (gray, left) and Beacon (blue, right) conditions. Each transparent colored circle represents one estimation by a participant and is linked to the corresponding POI flag by a transparent line, which represents the amount of error (Per POI and group views are available in the Supplementary). (B) The amount of error between estimated and actual POI locations for all groups and conditions. (C) Heat map of the routes walked in the Turn-by-turn (gray, left) and Beacon (blue, right) conditions. (D) The distance (decimal degrees) walked during the scavenger hunt of all groups and conditions. All data are presented as ± SEM, * p < 0.05

### There was no difference in the amount of distance traveled between groups or conditions

While all participants in the Turn-by-turn groups used the same route to visit all five POIs, the participants in the Beacon group were free to use any route (Figure 3C). Despite the differences in route, there was no difference in the total distance that was traversed by all groups (All groups: Mean = 2267.51 meters, SD = 245.93). A 2×2 ANOVA across group (Expert and Naïve) and condition (Turn-by-turn and Beacon) revealed no significant main effect of group (Figure 3D; F(1,49) = 1.45, p = 0.23), condition (F(1,49) = 0.52, p = 0.47), or interaction (F(1,49) = 0.05, p = 0.81).

## Discussion

The goal of the study presented here was to present an alternative to turn-by-turn navigation that is effective and is still engaging users into their environment to promote spatial learning. We show that using auditory beacons to navigate can lead to greater explorative behavior and the formation of more accurate mental maps of the surrounding environment when compared to turn-by-turn navigation. Thus, demonstrating that it is possible to use GPS technology and promote learning through active navigation. In this study, groups of both familiar (Experts) and unfamiliar (Naïve) participants were asked to find the locations of different POIs on a scavenger hunt across the Microsoft campus. Half of the groups were guided to the POIs using traditional turn-by-turn directions (Turn-by-turn) and the other half used the auditory navigation app Soundscape (Beacon) to find the POIs. While the pre-post performance showed little differences between the conditions in the Expert group, the Naïve Beacon group demonstrated a more accurate spatial representation of the Microsoft campus as reflected by the significant improvement (pre-post) in their ability to point to each of the POI locations, both from a familiar and a novel location, as well as to identify the POI locations on a simple overhead map of the area. It was not surprising that experts, who were familiar with the campus, did not exhibit a significant difference in learning as we expected them to have a reasonable mental map already formed.

There are several limitations of the current study that we feel are important to acknowledge. First, while we were able to test four different groups, we ideally would have had greater numbers in each of the individual groups. Time was a limiting factor, especially in our naïve summer interns as they were extremely busy throughout the day and the duration of their internship was short. Participants were often run during an extended lunch break or before/after work hours. Second, the Turn-by-turn groups were given verbal directions by an experimenter as opposed to a navigation app. One issue we encountered when testing turn-by-turn navigation apps was that these apps could not accurately inform a walking participant of when they needed to turn. The area we used for our study required participants to walk along paved and dirt paths within the Microsoft campus and the turn-by-turn navigation apps failed to reliably direct participants to each of the POIs. Lastly, this experiment was designed for walking navigation and the area chosen, containing complex intersections and buildings, was selected based on the absence of roads for safety concerns. Many turn-by-turn systems might be better suited for actual road or car navigation as the goal is less about learning the environment and more about getting to the desired location in a timely manner (additional details of how many turns and directions were given to participants can be found in the supplementary materials). We believe however that our results would also translate to a car driving situation, since turn-by-turn has been shown to reduce the ability to recall and sketch the maps in drivers (35), as well as pedestrians (36).

Nevertheless, this does not detract from the main point and results of this paper: (i) Turn-by-turn directions are a passive form of navigation that do not support spatial learning of the surrounding environment. (ii) By taking a more active role in their own navigation, participants were able to construct a better understanding of the spatial layout of a real-world environment. Using a spatial auditory navigation app to promote active navigation, we can theoretically engage the hippocampus (7, 27, 37–40) resulting in a stronger and more accurate mental map of the explored space.

It is clear that GPS technology is an integral part of our everyday lives and it is unlikely that society would give up such an important technological advance. While the benefits of using this technology are undeniable, it is important to consider how navigation should be implemented, as GPS is a global positioning system that can support different forms of navigation. Rather than replacing the decision-making process, our work proposes to augment navigation with a rich sensory experience that engages perception and cognitive processing. After all, just as technology cannot replace the benefits of physical exercise, it cannot replace the benefits of staying mentally active.

While our ability to navigate might seem like an accessory to our modern lives, at one point these navigational skills were critical for our own ancestral survival. It is not surprising that some of the most powerful mnemonic techniques, for example the memory palace, involve setting mental pictures of items or facts in locations in an imaginary place, such as a building or a town.

Memories become easier to recall when they’re associated with physical locations, even if only in the imagination (41,42). Nor it is surprising that some of the first signs of cognitive decline associated with Alzheimer is the deterioration of spatial awareness (23). Importantly, recent studies have found that exercising spatial cognition might protect against age-related memory decline (19). Therefore, it is important that, while using technology, we still engage with the world around us as we know it is important for brain health.

In this paper we are not suggesting that we need to outright reject technology, we simply need to rethink the way we interact with technology, to design and create in ways that engage the brain rather than ignore it. Auditory beacon navigation is a step into a new era in which automation does not simply replace our evolutionary functions and remove us from our environment, but rather feeds into our sensory inputs to promote active engagement with the world around us. In that regard, GPS navigation based on auditory beacons is a sensory augmentation that helps us create a stronger connection with our environment.

## Acknowledgments

The authors would like to thank Microsoft Research for the continued support to the research as well as the Enable team who created Microsoft Soundscape: Adam Glass, Melanie Kneisel, Daniel Tsirulnikov, Amy Karlson, Jarnail Chudge, Arturo Toledo, Emily Greene, Rico Malvar. The authors would also like to thank Matthew Bennett and Scott Reitherman for the interoceptive sound design of the Beacons. And the members of the Extended Perception Interaction and Cognition (EPIC) Team: Ken Hinckley, Bill Buxton, Eyal Ofek, and Ability team: Merrie Morris and Ed Cutrell. Finally, also thank the Stark lab at University of California Irvine.

## Author Contributions

GDC and MGF wrote the manuscript. AM designed the beacon functioning and soundscape app. GDC designed the experiment. AF built the scavenger hunt on the soundscape app. GDC analyzed the data and MGF did the heatmap visualizations on GPS data. GDC and MGF did the data collection and recording. All authors edited the manuscript.

## Other supplementary materials for this manuscript include the following

**Fig. S1.**
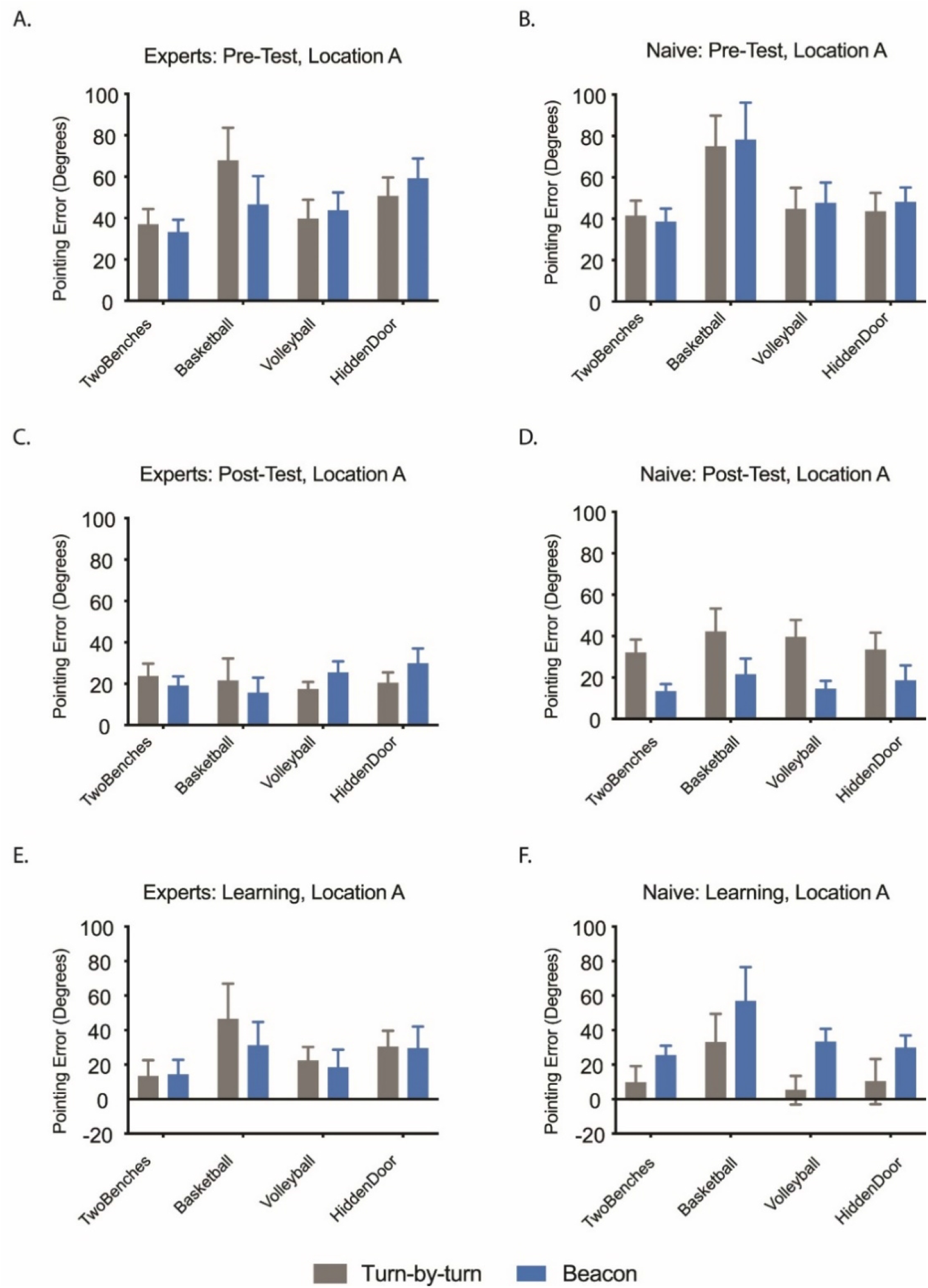
Individual POI data of both groups from Location A. (A) Pre-Test pointing error of Experts for all individual POIs from Location A. (B) Pre-Test pointing error of Naïve for all individual POIs from Location A. (C) Post-test pointing error of Experts for all individual POIs from Location A. (D) Post-Test pointing error of Naïve for all individual POIs from Location A. (E) The improvement in pointing from Pre-Test to Post-Test of Experts from Location A. (D) The improvement in pointing from Pre-Test to Post-Test of Naïve from Location A.

**Fig. S2.**
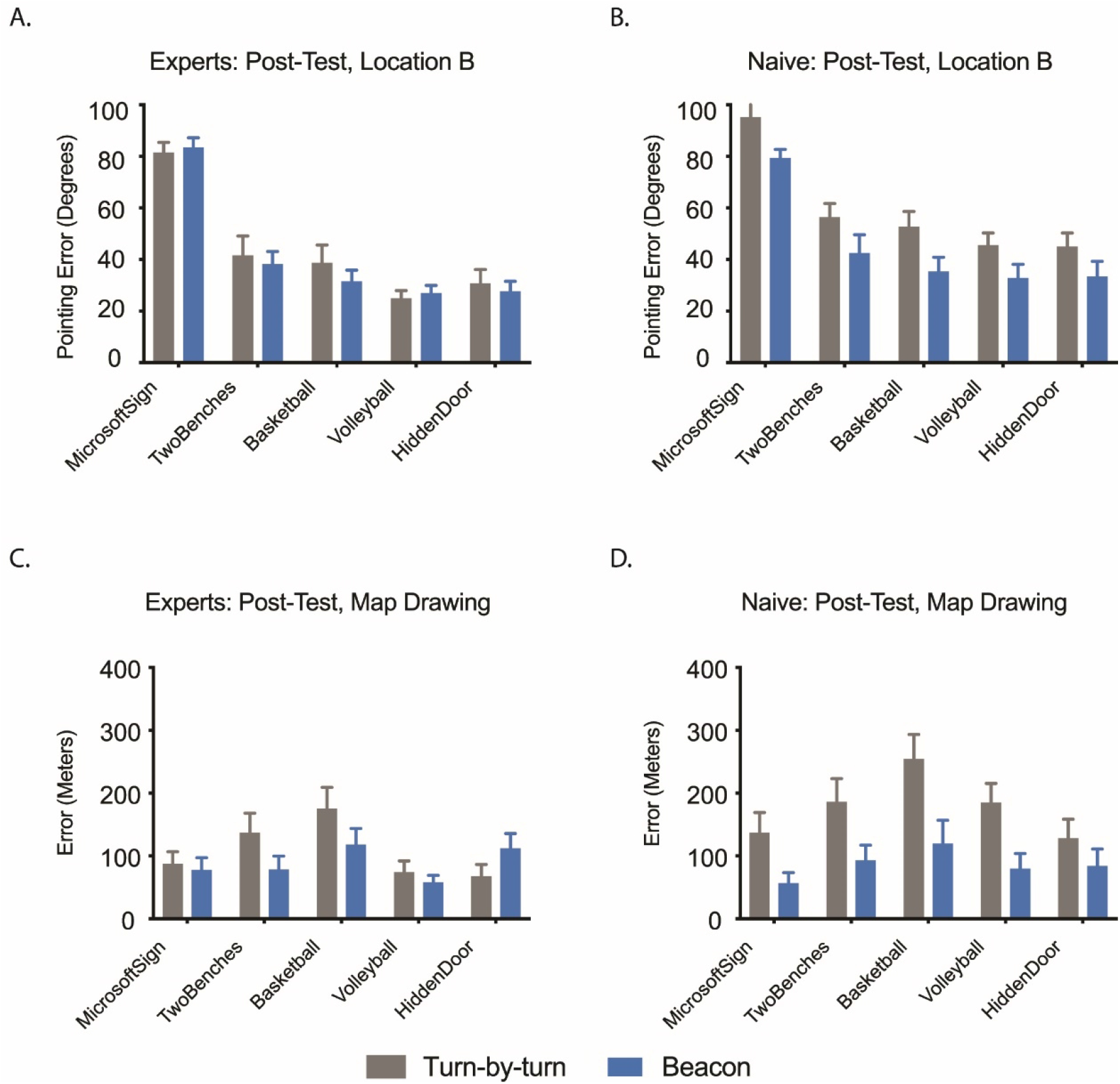
Individual POI data of both groups from Location B and Map Drawing. (A) Post-Test pointing error of Experts for all individual POIs from Location B. (B) Post-Test pointing error of Naïve for all individual POIs from Location B. (C) Map Drawing performance on all individual POIs for Experts. (D) Map Drawing performance on all individual POIs for Naïve.

**Fig. S3.**
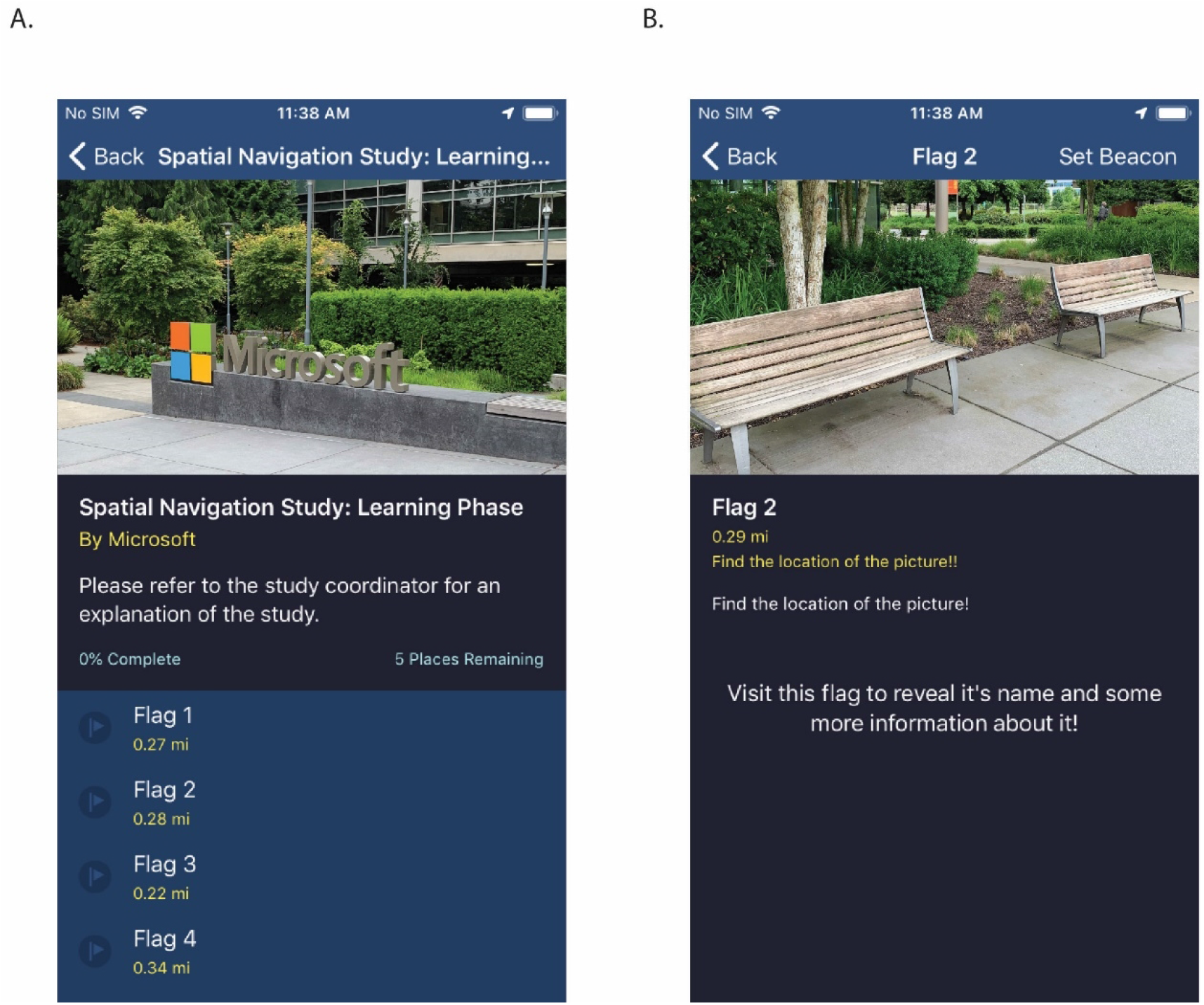
Example images of the Soundscape app scavenger hunt. (A) Overview of the scavenger hunt and picture of the first POI. (B) After a POI is found, an image of the next one shows up.

**Fig. S4.**
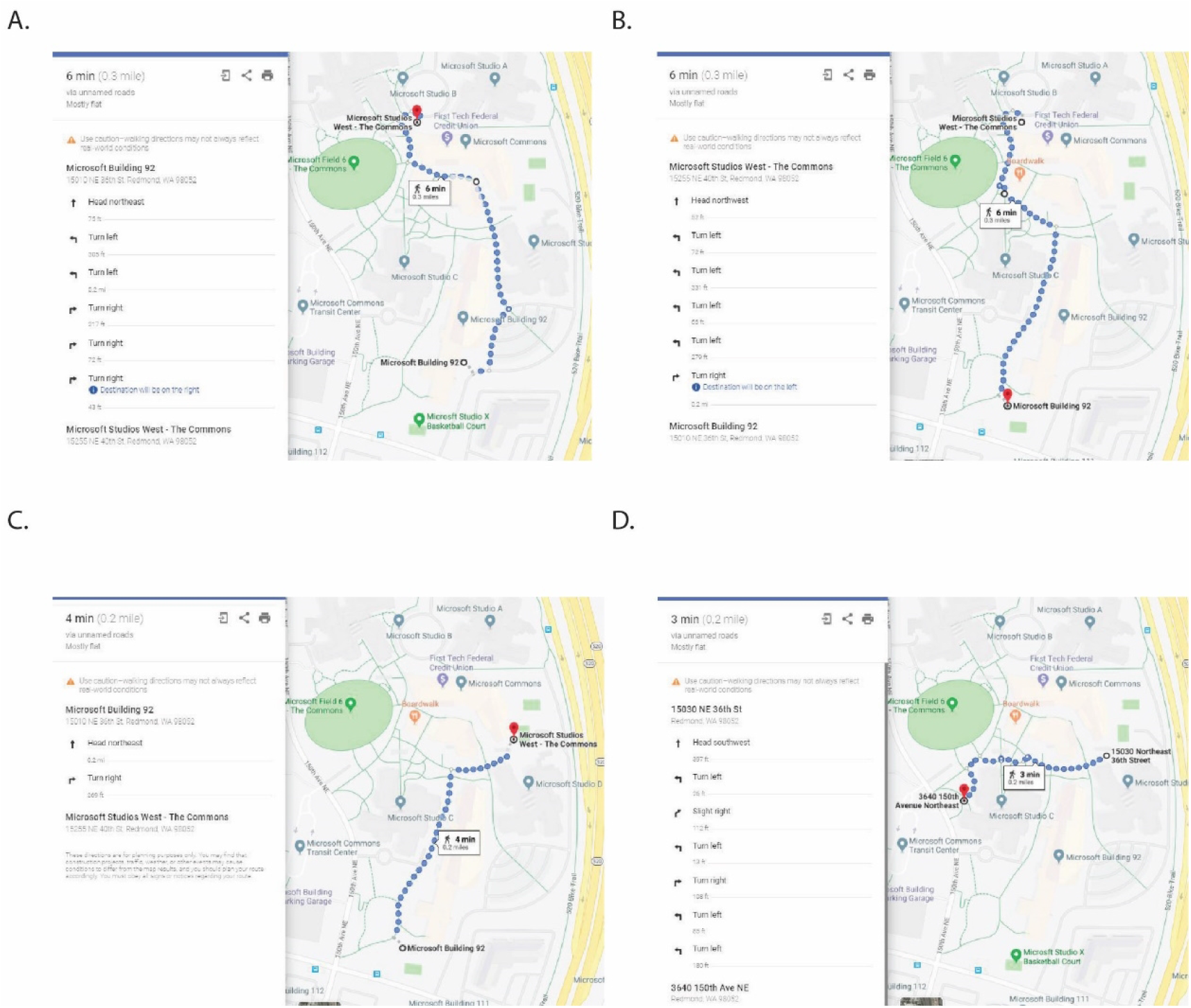
Directions used for the Turn-by-turn group. (A) Directions from Microsoft Sign to Two Benches. (B) Directions from Two Benches to Basketball Court. (C) Directions from Basketball Court to Volleyball Court. (D) Directions from Volleyball Court to Hidden Door.

**Fig. S5.**
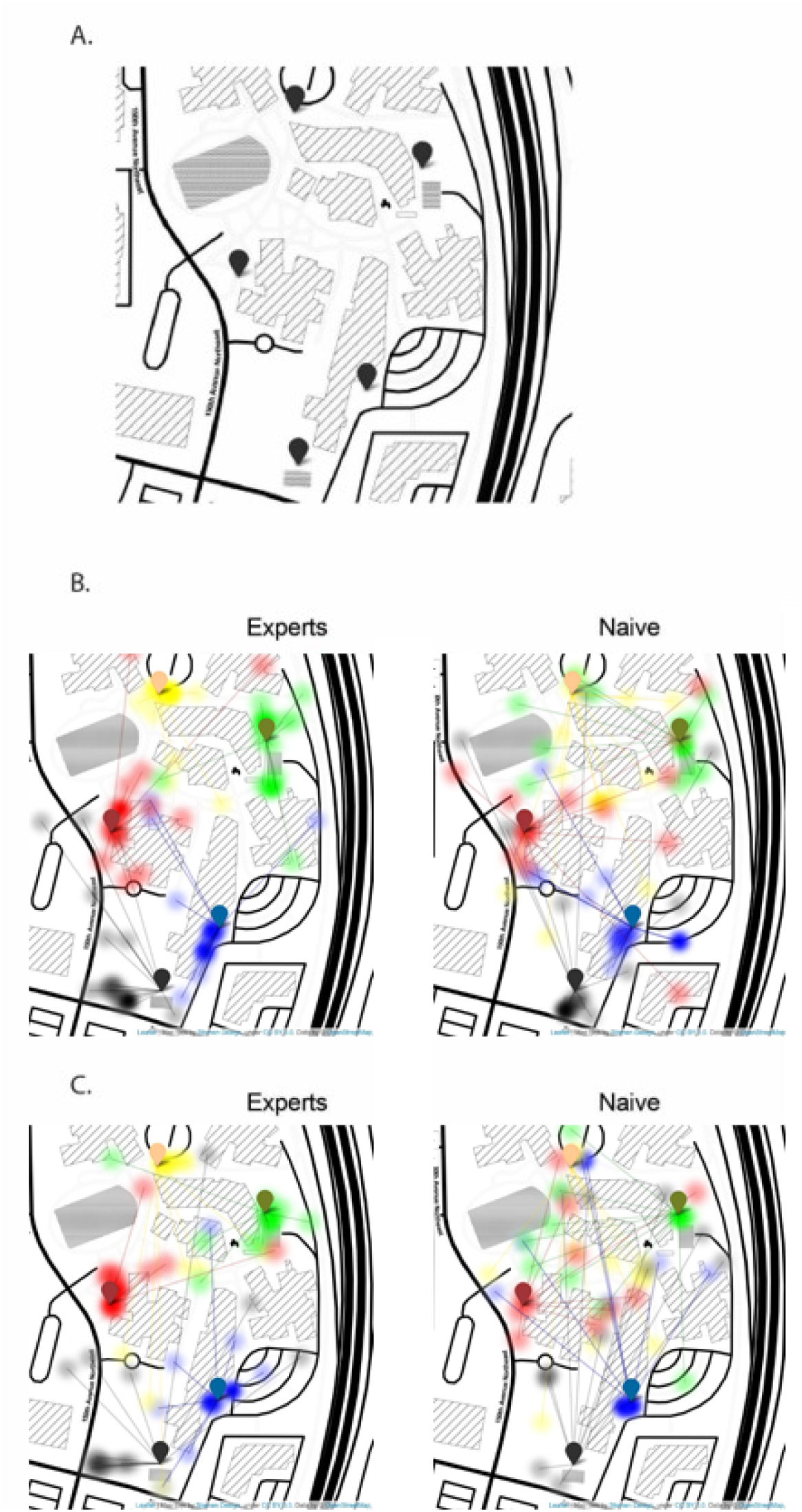
Map Drawing task and group performance. (A) Simple map with the marked destinations of the scavenger hunt. (B) Performance of both groups in the Beacon group. (C) Performance of both groups in the Turn-by-turn group.

**Fig. S6.**
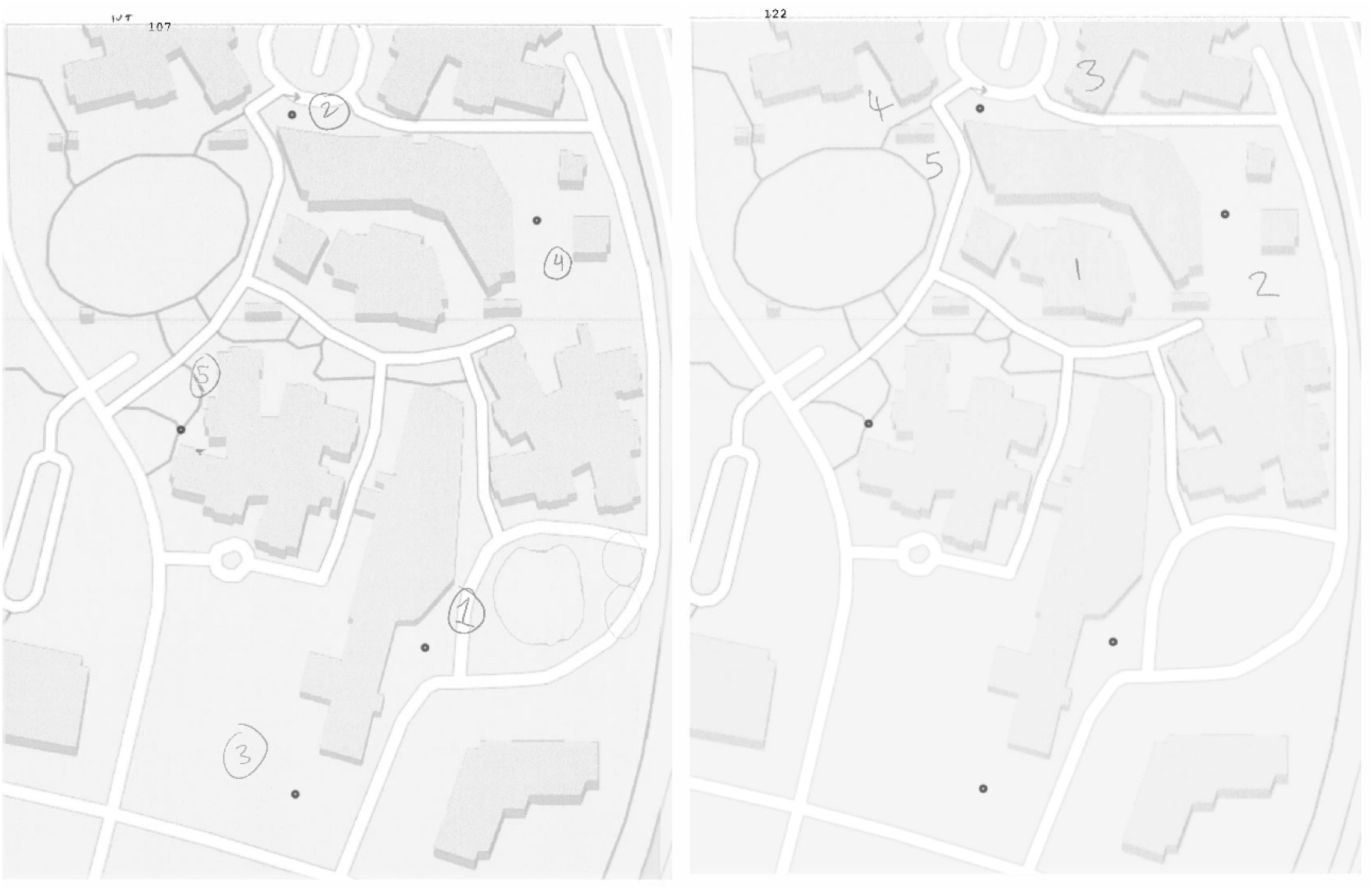
Real maps with user handwritten numeric locations (numbers of landmarks: 1,2,3,4,5) on the Map Drawing task. For illustration, dots are placed at the scavenger hunt latitude and longitude targets. Left) Drawn map of a participant exhibiting a high-quality mental map, with minimal error between real landmarks and their mental map. Right) The map drawn by a participant who clearly had not developed an accurate mental map after the navigation.

**Fig. S7.**
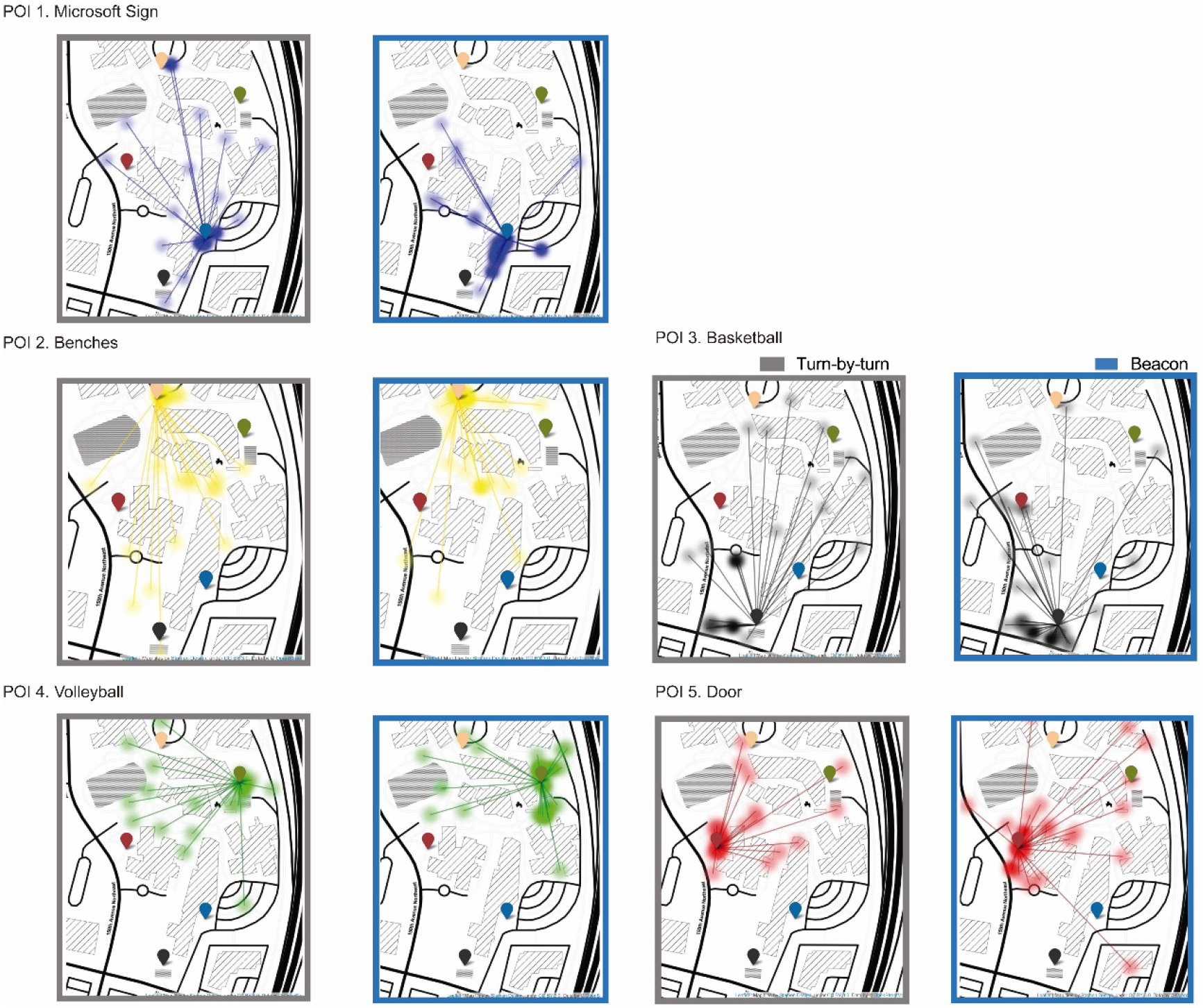
Map Drawing task and group performance for each POI destinations of the scavenger hunt. For the Turn-by-Turn and Beacon conditions.

**Movie S1 (separate file).** The video contains an explanation of the results, footage of the participants going through the scavenger hunt and audio of the actual GPS app.

## Footnotes

https://www.microsoft.com/en-us/research/product/soundscape/

